# New *Aequorea* fluorescent proteins for cell-free bioengineering

**DOI:** 10.1101/2022.12.08.519681

**Authors:** Christopher Deich, Nathaniel J. Gaut, Wakana Sato, Aaron E. Engelhart, Katarzyna P. Adamala

## Abstract

Recently, a new subset of fluorescent proteins has been identified from the Aequorea species of jellyfish. These fluorescent proteins were characterized *in vivo*; however, there has not been validation of these proteins within cell-free systems. Cell-free systems and technology development is a rapidly expanding field, encompassing foundational research, synthetic cells, bioengineering, biomanufacturing and drug development. Cell-free systems rely heavily on fluorescent proteins as reporters. Here we characterize and validate this new set of Aequorea proteins for use in a variety of cell-free and synthetic cell expression platforms.

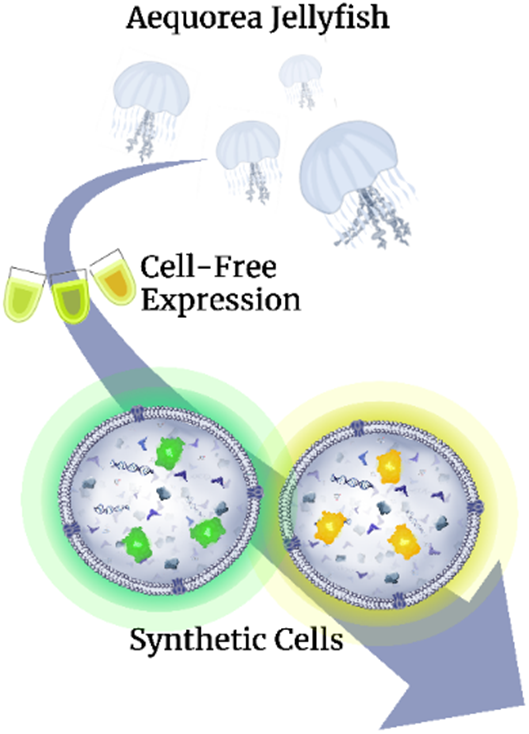

## Introduction

Cell-free protein expression systems have become a cornerstone of synthetic biology^1–4^, offering the flexibility of in vitro systems while maintaining some of the life-like complexity of live biological systems. Encapsulating these systems within liposomes allows for the construction of synthetic minimal cells, a technology with far reaching uses as a model system and technology development platform^5–7^. To the same extent as live biological systems, fluorescent proteins (FPs) as reporters are incredibly useful for monitoring gene expression to better optimize and develop novel assays within these synthetic systems^8–10^.

Recently, a new suite of FPs has been discovered within the species of jellyfish *Aequorea* cf. *australis*^11^. Because these proteins have not been validating within any cell-free expression (CFPE) platforms, we investigated their efficacy within the *E. coli*-based cell-free protein expression system, TxTl (short for transcription-translation protein synthesis), perhaps the most commonly used and user-friendly CFPE system. We further characterize these new FPs within liposomal bioreactors, encapsulating TxTl, and tested efficacy within commercially available CFPE systems from a variety of species.

In all, we showcase the ability of these new FPs to be used as synthetic biology tools within cell-free systems. Virtually all cell-free systems rely heavily on FP technology and adding more FPs to the synthetic biology color palette will allow for more options to be available when FPs outside of the commonly available ones are needed.

## Results and discussion

A suite of fluorescent proteins genes from *Aequorea* were received with each protein under the control of the T7 promoter, a promoter common in cell-free systems^12,13^. To increase expression levels, we swapped this promoter for the T7Max promoter, shown to significantly increase expression levels in TxTl^14^ and also swapped the original UTR regions for ones optimized for cell free expression shown in previous work^15^. In our pilot tests, several proteins did not express well under any tested circumstances (data not shown); however, three in particular gained marked increases in expression levels (**Fig yS4**) and were the ones carried forward for further study. These three were termed mAbb and tdAbb, monomer and tandem dimer green fluorescent proteins (GFP) respectively, as well as mAbb_Y, a monomer yellow fluorescent protein (YFP) variant. For all following characterization assays, these fluorescent proteins were compared to classical GFP for mAbb and tdAbb and YFP for mAbb_y.

After using a TxTl reaction to express these proteins, we demonstrate that all three of these proteins have a similar maturation time to GFP and YFP, reaching 80-90% of peak expression levels within the first 2-3 hours (**Fig 1a-e**), indicating that they would have similar viability as reporters when this maturation time length is desired. tdAbb reaches comparable peak fluorescence as GFP, while the two monomers have marked lower peak fluorescence compared to their respective counterparts; however, the levels are still within 80% of GFP and YFP, validating them as viable reporters for cell-free systems. The proteins’ expression levels were validated with a Western blot (**Fig 1f**), supporting the trends discussed earlier. Interestingly, tdAbb displays a weak lower-molecular weight band, indicating that a fraction of the protein expressed as a monomer.

**Figure 1.**
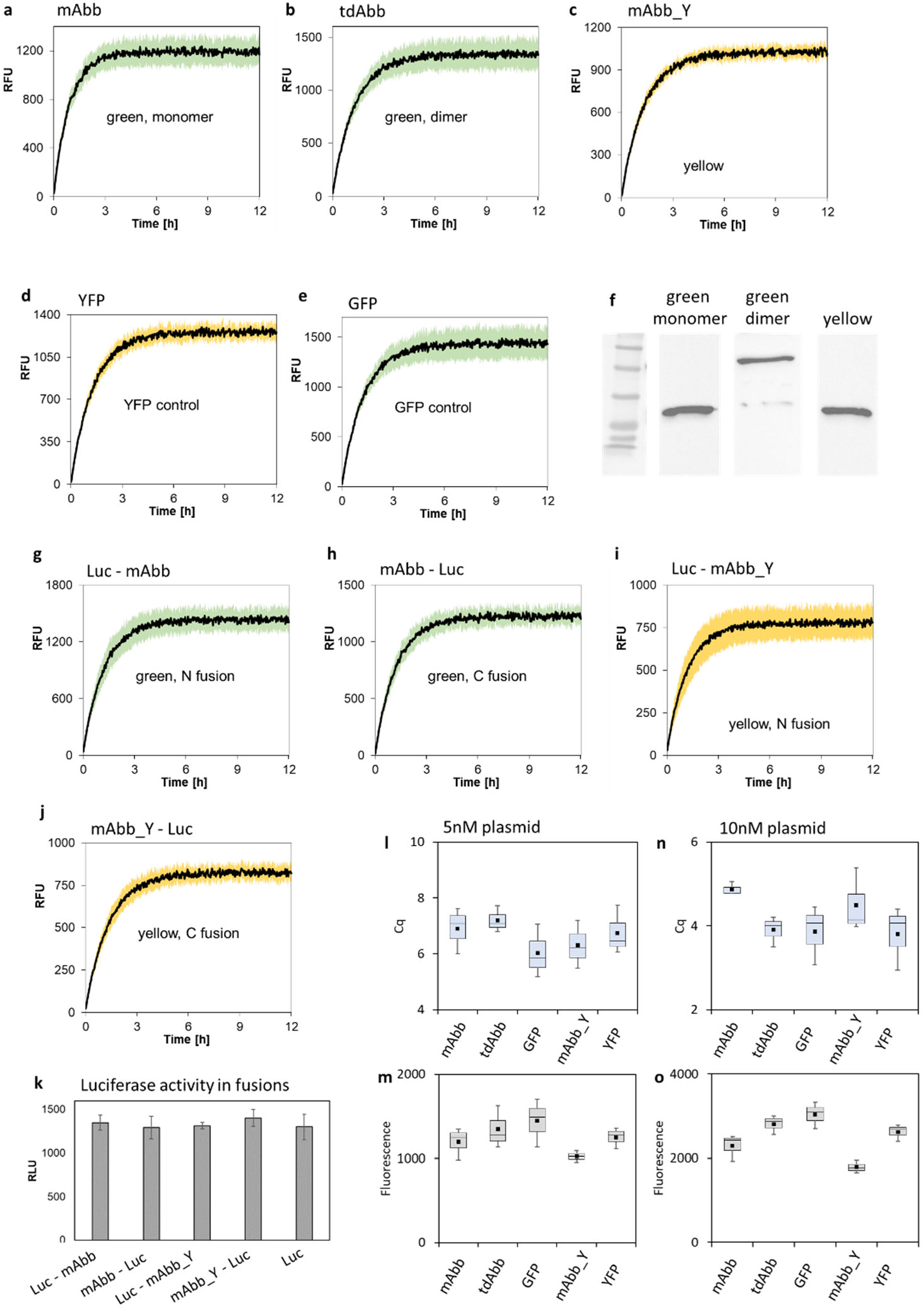
The new *Aequorea* fluorescent proteins in bacterial cell-free expression system. **a-e**: Time course of expression of green monomer (mAbb, panel **a**), green dimer (tdAbb, panel **b**), green control (GFP, panel **e**), yellow monomer (mAbb_Y, panel **c**) and yellow control (YFP, panel **d**) in *E. coli* TxTl. **f**: Western Blot analysis of expression of the green monomer, dimer and yellow protein. Original unprocessed gels are on **Figure yS1, yS2** and **yS3**. **g-k**: Performance of the new fluorescent proteins in N-/C-terminal fusions with Firefly luciferase via measurements of the fluorescent protein expression and luciferase activity for each fusion. The fluorescence expression time course was measured for green monomer N- (Luc - mAbb, panel **g**) and C-terminal fusion (mAbb - Luc, panel **h**), and yellow monomer N- (Luc – mAbb_Y, panel **i**) and C-terminal fusion (mAbb - Luc, panel **j**), followed by a luciferase activity assay for each of those fusion proteins (panel **k**, with luciferase without fluorescent protein as control). **l-o**: The fluorescence and mRNA abundance of the fluorescent proteins at two different plasmid concentrations. Fluorescence of the green monomer (mAbb), green dimer (tdAbb), green control (GFP), yellow monomer (mAbb_Y) and yellow control (YFP) was measured in TxTl reactions with 5nM (panel **l**) and with 10nM (panel **n**) of each plasmid, followed by RT qPCR analysis of mRNA abundance (panel **m** for 5nM plasmid and panel **o** for 10nM plasmid). All experiments were performed in triplicate. Shaded areas on panels **a-e** and **g-j**, and error bars on panel k indicate SEM.

Often, fluorescent proteins are fused with another protein to monitor the expression levels of this protein of interest^16,17^. To test the viability of these new fluorescent proteins in such a system, we cloned a fusion protein consisting of luciferase, a commonly used and well-characterized protein within the synthetic biology field^18–20^, to either the N- or C- termini of the monomers FPs (mAbb or mAbb_y). Monitoring expression of these fusion proteins in a TxTl reaction revealed no decrease in fluorescence regardless of the termini fused (**Fig 1g-j**) as well as no decrease in luciferase activity (**Fig 1k**).

To investigate the lower fluorescence signal displayed by the monomeric FPs, we assayed their mRNA levels post-TxTl via qPCR. All samples had similar mRNA expression levels with 5nM of input plasmid (**Fig 1l,m**), indicating that the monomeric FPs are intrinsically less fluorescent than the dimer or GFP and YFP. This trend held true when we varied the plasmid concentration of the FPs to 10nM; additionally, the expression output scaled at the same rate as GFP and YFP (**Fig 1n,o**). This was a promising result, since it is very important in cell-free systems to be able to scale gene expression with template concentration, allowing for the expression stoichiometric protein ratios^21^.

Because cell-free systems are often encapsulated within liposomal membranes to construct minimal synthetic cells^22–25^, we proceeded to test the viability of these proteins to function well in such a system. We encapsulated the above TxTl expression systems within POPC/cholesterol membranes and imaged the liposomes after a 12-hour incubation (**Fig 2a-e**) and further quantified the fluorescence of individual liposomes (**Fig2f**). All new FPs had significant fluorescence visible by eye and the relative fluorescence followed the previous observed trend, the monomer FPs yielding around 80-90% the fluorescence of GFP and YFP, while tdAbb was comparable to GFP. To ensure these FPs did not destabilize the liposomal membranes, we encapsulated calcein alongside the TxTl reaction and assayed for membrane leakage by running the liposomes through a size-exclusion column. In all samples, less than 2% total calcein was detected in the low-molecular weight fraction (**Fig 2g**), indicating liposome membrane maintained integrity. Overall, these new FPs expressed at similar levels when encapsulated within liposomes and do not cause any membrane destabilization.

**Figure 2.**
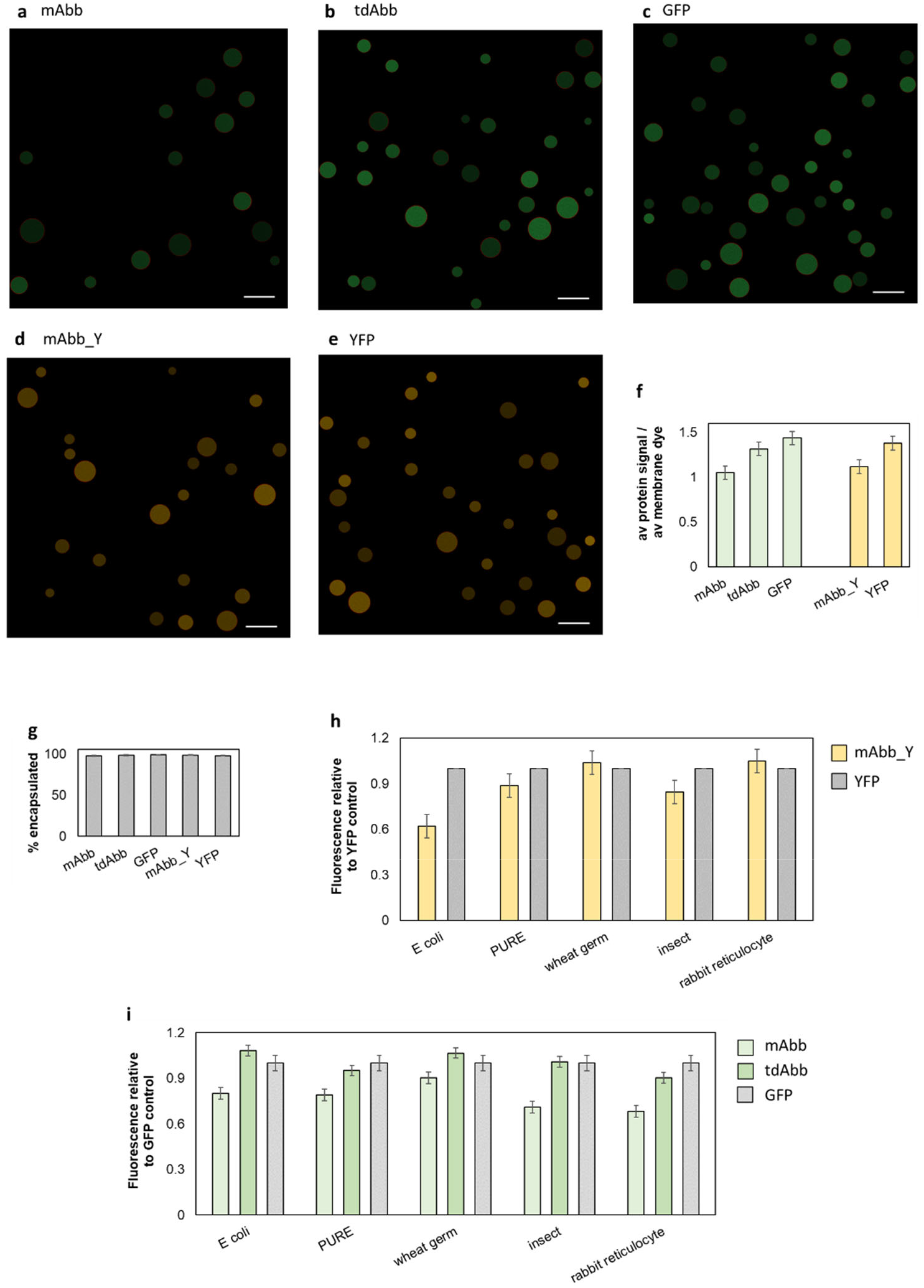
The new fluorescent proteins in different contexts: in synthetic cells, and in a variety of cell-free expression systems. **a-e**: New fluorescent proteins, *E. coli* TxTl, and T7 RNA polymerase were encapsulated within red-labelled (via Rhodamine-PE) POPC/cholesterol liposomes. These synthetic cells were then imaged under the fluorescent microscope after a 12-hour incubation. The green monomer mAbb is on panel **a**, green dimer tdAbb is on panel **b**, green control GFP is on panel **c**, yellow monomer mAbb_Y is on panel **d** and yellow control YFP is on panel **e.** **f**: Average fluorescence in the green and yellow channels was quantified for 10 representative synthetic cell microscopy images from each sample, each with three independent replicates, then normalized the fluorescent protein signal to the red channel signal (the membrane dye signal, quantifying the total liposome density). **g**: The membrane stability of synthetic cells expressing the new fluorescent proteins was assayed via leakage tests. Calcein was encapsulated in synthetic cells expressing each of the fluorescent proteins. After a 12-hour incubation, calcein leakage was measured using size exclusion chromatography. Representative individual purification traces are on **figure yS5**. **h** and **i**: Expression of the green monomer (mAbb, panel **i**), green dimer (tdAbb, panel **i**), green control (GFP, panel **i**), yellow monomer (mAbb_Y, panel **h**) and yellow control (YFP, panel **h**) in different cell-free translation systems. The expression was tested using home-made *E. coli* TxTl and commercial kits for PURE, wheat germ extract, insect *Spodoptera frugiperda* Sf21 cell line extract and rabbit reticulocyte extract. All experiments were performed in triplicate. Error bars indicate SEM.

Lastly, we tested the versatility of these new FPs by comparing their expression to GFP and YFP in the following in vitro expression commercial systems, following the manufacturer’s directions: : PURE system composed of *E. coli* translation machinery purified individually^26^, wheat germ extract^27^, *Leishmania tarentolae* extract^28^, insect *Spodoptera frugiperda* Sf21 cell line extract^29^, and rabbit reticulocyte extract^30^. Due to expression variability in each system, the results were normalized the GFP for mAbb and tdAbb; and YFP for mAbb_Y. The earlier trends also held true here, with tdABB fluorescence comparable to GFP and mAbb and mAbb_Y expressing slightly lower than GFP and YFP (**Fig 2h-i**). Those results imply that the new fluorescent proteins are just as versatile as the classical fluorescent proteins of GFP and YFP, allowing them to be used in a wide variety of cell-free systems.

Overall, we demonstrate the practicality of these new Aequorea FPs when used in cell-free expression systems. Specifically, we characterized their expression and fluorescence profiles, ability to function within a liposomal environment, and versatility within a host of commonly-used cell-free expression kits. We are proud to offer our data to give confidence for the community to make use of these new FPs within a variety of synthetic biology applications. We also believe it is important to continue characterizing live cell technologies within cell-free systems, since, in our case, the results translated favorably over to an in vitro environment.

## Acknowledgments

We thank Dr. Nathan Shaner for the gift of plasmids containing the new fluorescent proteins tested in this work. This work was supported by, NSF award 1807461 SeMiSynBio Very Large scale genetic circuit design and automation, NSF award 1840301 RoL:FELS:RAISE Building and Modeling Synthetic Bacterial Cells, NSF award 2123465 Synthetic P-bodies: Coupling gene expression and ribonucleoprotein granules in synthetic cell vesicles for sensing and response. Wakana Sato was supported by the Funai Overseas Scholarship of The Funai Foundation for Information Technology.

## Materials and methods

### Cloning FPs under T7max promoter

pCI-T7Max-UTR1-CTerminus8xHis-T500 (Addgene, 178419) with a carbenicillin-resistant gene was used as the plasmid backbone. The plasmid was digested with SpeI-HF (NEB, R3133L) and MluI-HF (NEB, R3198L), cutting downstream of the T7Max promoter. PCR primers for fluorescent protein genes were designed to add SpeI and MluI restriction cut sites at the N- and C-terminus of the genes, respectively. The genes were PCR amplified using Phusion polymerase (NEB, M0530L), followed by 1% agarose gel electrophoresis and gel purification using a gel extraction kit (Epoch Life Science, 2260050). Recovered PCR products were digested with SpeI-HF and MluI-HF. The digested backbone and gene inserts were isolated using a PCR clean-up kit (Epoch Life Science, 2360250). The backbone was treated with rSAP (NEB, M0371L) to prevent self-ligation of the backbone.

The insert and backbone were ligated with T4 DNA ligase (NEB, M0202L). The ligated plasmid was transformed into *E. coli* BL21_DE3 and plated on LB agar plates with carbenicillin. Multiple clones were grown in LB liquid cultures, and plasmid DNA was isolated using a mini-prep kit (Epoch Life Science, 2160250). The DNA sequence was analyzed using a primer that binds upstream of the T7Max promoter and confirmed the transition into ORF.

### Western Blot Analysis of T7max FPs

Triplicated samples from the brightness comparison (**fig 1a-c**) were fractionated on 10% SDS-Page gel. The gels were transferred to a 0.2 μ m nitrocellulose membrane at 100V for 1 hour in cooled transfer buffer (25mM Tris, 192mM Glycine, 10% methanol). The membrane was incubated with 5% nonfat milk in TBST for 1 hour followed by the addition of primary antibodies (BioLegend, 652505). After 1 hour of the primary antibody incubation, the membrane was rinsed three times with TBST followed by three 10 minutes washes in TBST. Secondary antibodies (BioLegend, 405306) were added to 5% nofat milk in TBST and incubate for 1 hour. After the incubation, the membrane was rinsed three times with TBST followed by three 10 minutes washes in TBST. Blots were developed with SuperSignal (Thermo Scientific) immunoblotting detection system according to manufacturer’s protocols. Blots were imaged using the ChemiDoc MP Imaging System (BioRad) running Image Lab version 5.2.1. The high resolution chemiluminescent image and colorimetric image were merged with GelQuantNet software version 1.8.2 before the analysis.

### Bacterial TxTl

TxTl protocol was adapted from the Noireaux and Jewett protocols^31,32^. 3x cell-free prep was prepared using Rosetta 2 (Novagen, 71400), following the method described previously^24^. TxTl reaction is composed of the following: 12 mM Magnesium glutamate; 140 mM potassium glutamate; 1 mM DTT; 1.5 μM T7 RNA polymerase; 0.4 U/µl Murine RNase Inhibitor (NEB, M0314S); 1x cell-free prep; 1x energy mix; and 1x amino acid mix.

10x energy mix composition was the following: 500 mM HEPES, pH 8; 15 mM ATP; 15 mM GTP; 9 mM CTP; 9 mM UTP; 2 mg/mL E. coli tRNA; 0.68 mM Folinic Acid; 3.3 mM NAD; 2.6 mM Coenzyme-A; 15 mM Spermidine; 40 mM Sodium Oxalate; 7.5 mM cAMP; 300 mM 3-PGA.

10x amino acid mix was prepared by mixing 20 mM of the following amino acids: alanine, arginine, asparagine, aspartic acid, cysteine, glutamic acid, glutamine, glycine, histidine, isoleucine, leucine, lysine, methionine, phenylalanine, proline, serine, threonine, tryptophan, tyrosine, and valine. Those amino acids were dissolved in pH 6.5, 400 mM potassium hydroxide solution.

The template plasmid concentrations were 10 nM, as well as no plasmid reactions for the negative control. Each reaction was triplicated. TxTl reactions were incubated at 30°C for 12 hours using SpectraMax plate readers. The fluorescence data was recorded at Ex 408/ Em 509 nm (EGFP) or Ex 505/ Em 527 nm (YFP). The TxTl samples were frozen after the 12 hours incubation for using in western blot analysis.

### Rt qPCR

The DNA in TXTL reaction was degraded with TURBO DNase (Invitrogen, AM2238). The proteins in the TXTL were denatured by incubating at 75°C for 15 minutes. The denatured proteins were removed by pelleting through centrifugation.

The reverse transcription reaction was performed by mixing the following: the DNase-treated sample, 1 μM primer, 10 mM DTT, 0.5 mM dNTP (Denville, CB4430-2), 5 U/μl Protoscript II reverse transcriptase (NEB, M0368X), 1x Protoscript II reverse transcriptase buffer, and 0.4 U/μl RNase Inhibitor.

The qPCR reaction was performed by mixing the following: the reverse-transcribed DNA, 0.8 μM FW and RV primers, 1x of OneTaq Hot Start 2X Master Mix with Standard Buffer (NEB, M0484L). The qPCR was performed on CFX96 Touch Real-Time PCR Detection System (BioRad). The amplification curves plotted through CFX Maestro Software to determine Cq values and averages across 3 replicates of each sample were calculated separately.

### Other extracts

The expression in extracts other than E. coli (described above), was performed using commercial kits. We used PURE system (NEB), wheat germ extract (Promega), insect *Spodoptera frugiperda* Sf21 cell line extract (Promega), and rabbit reticulocyte extract (Promega). All reactions were performed according to manufacturer’s instructions.

PURE expression plasmid was the same as used for E. coli extract. Wheat germ extract expression cassette was designed with UTR sequences based on Promega pF3 WG (BYDV) Flexi vector. *Spodoptera frugiperda* Sf21 cell line extract expression cassette was designed with UTR sequences based on Promega pF25K ICE T7 Flexi vector.

### Liposomes

Liposomes were prepared according to the DSCF protocol, using 3D printed device described before.^33^

Briefly, thin lipid films prepared by evaporating chloroform from mixture of lipids were dissolved in mineral oil. Microcapillary was fitted into the 3D printed microfluidic device. 100uL of the lipid oil solution was placed over 30uL of 100mM HEPES, 900mM Glucose, pH 7.5. The capillary was filled with the TxTl solution, and placed over the lipid oil buffer solution in a tube. The tube was centrifuged at 1,600 RCF for 3 minutes. Liposomes were recovered from the bottom of the tube after removing the capillary assembly.

## Supplementary Information

**Figure yS1.**
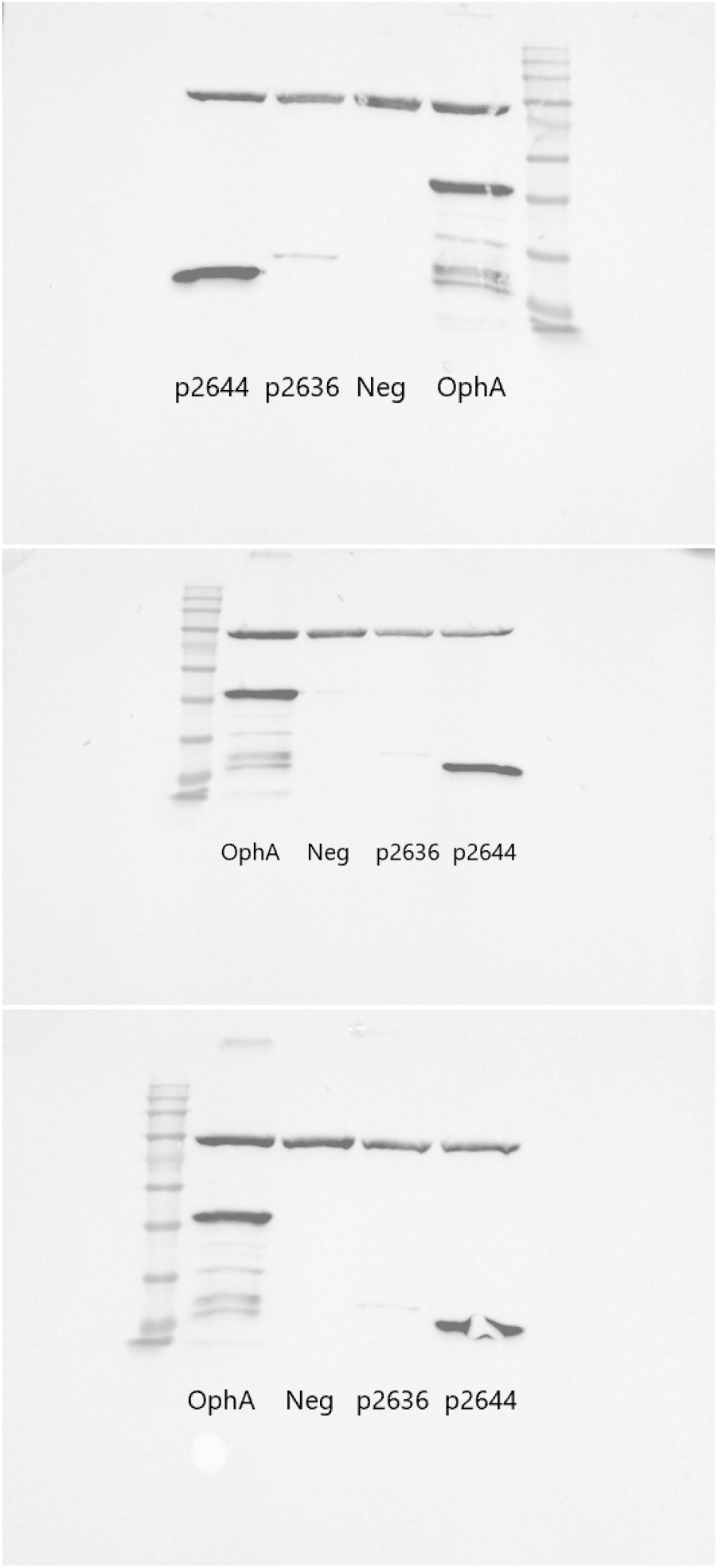
Western blot analysis of green monomer protein, sample *2644p*. The ladder is BLUEstain 2 Protein ladder.

**Figure yS2.**
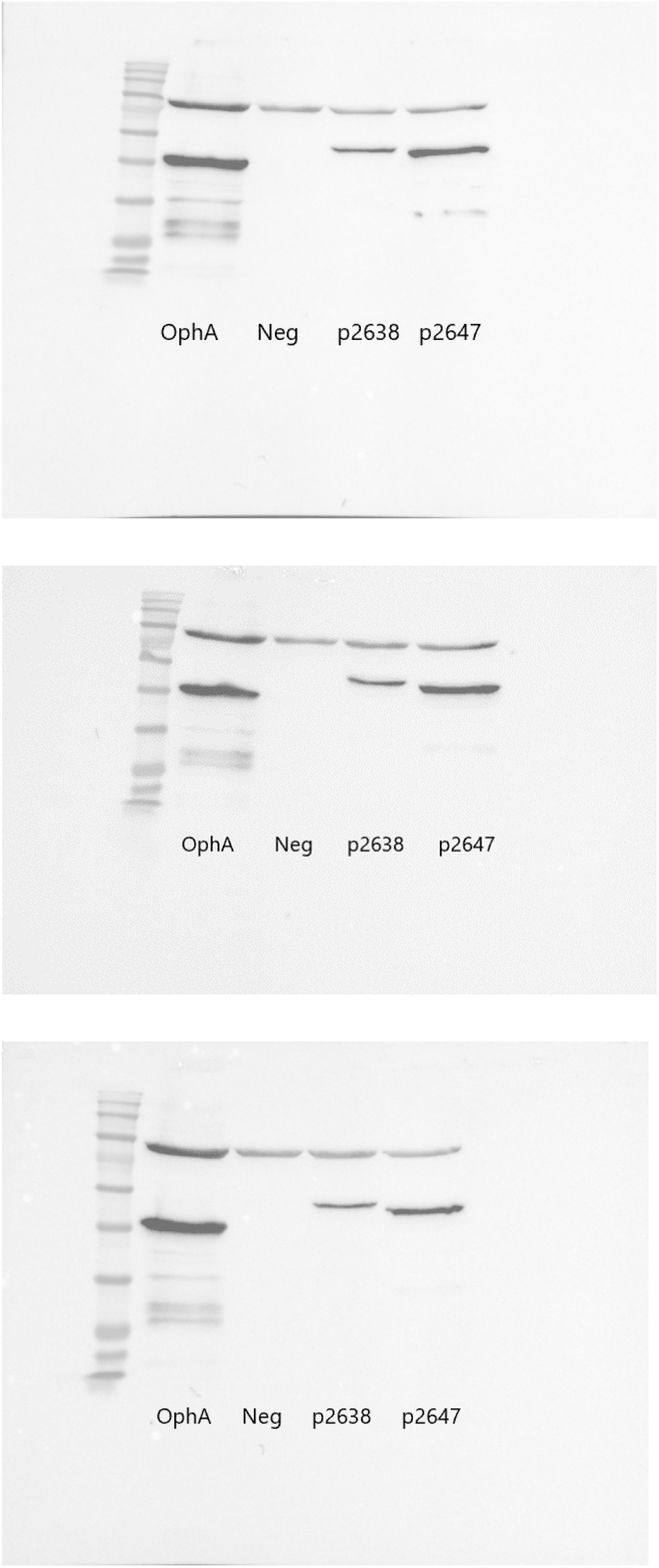
Western blot analysis of green dimer protein, sample *2647p*. The ladder is BLUEstain 2 Protein ladder.

**Figure yS3.**
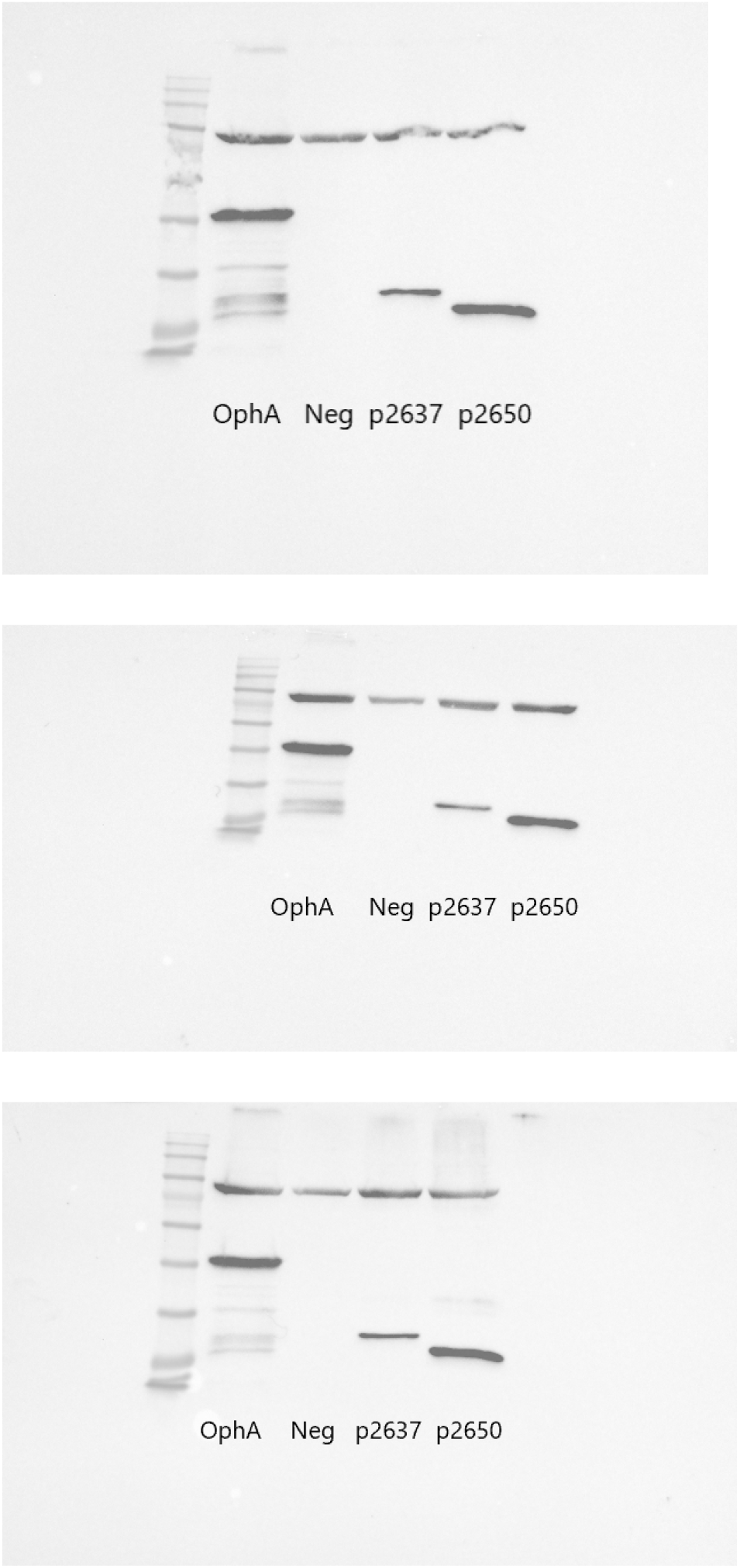
Western blot analysis of yellow protein, sample *2650p*. The ladder is BLUEstain 2 Protein ladder.

**Figure yS4.**
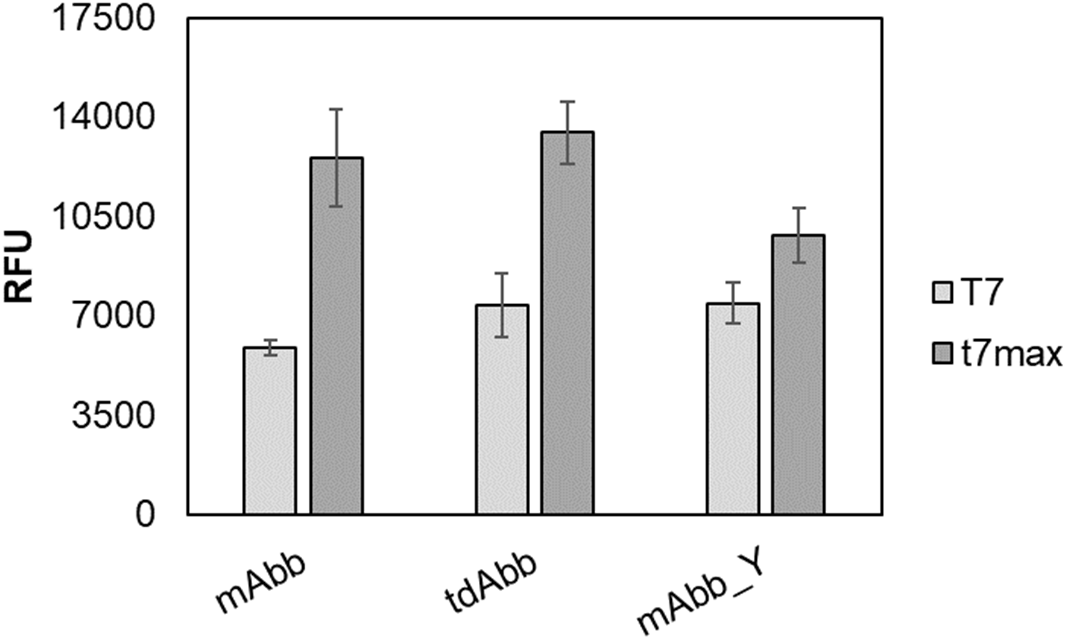
Comparison of expression of the new fluorescent proteins in original plasmids under T7 promoter, not optimized for cell-free expression, vs the same proteins in plasmids optimized for bacterial TxTl expression with T7 max promoter. Expression in E coli TxTl according to the protocol described in Materials and methods, 5nM plasmid in each rection. Error bars indicate SEM, n=3.

**Figure yS5.**
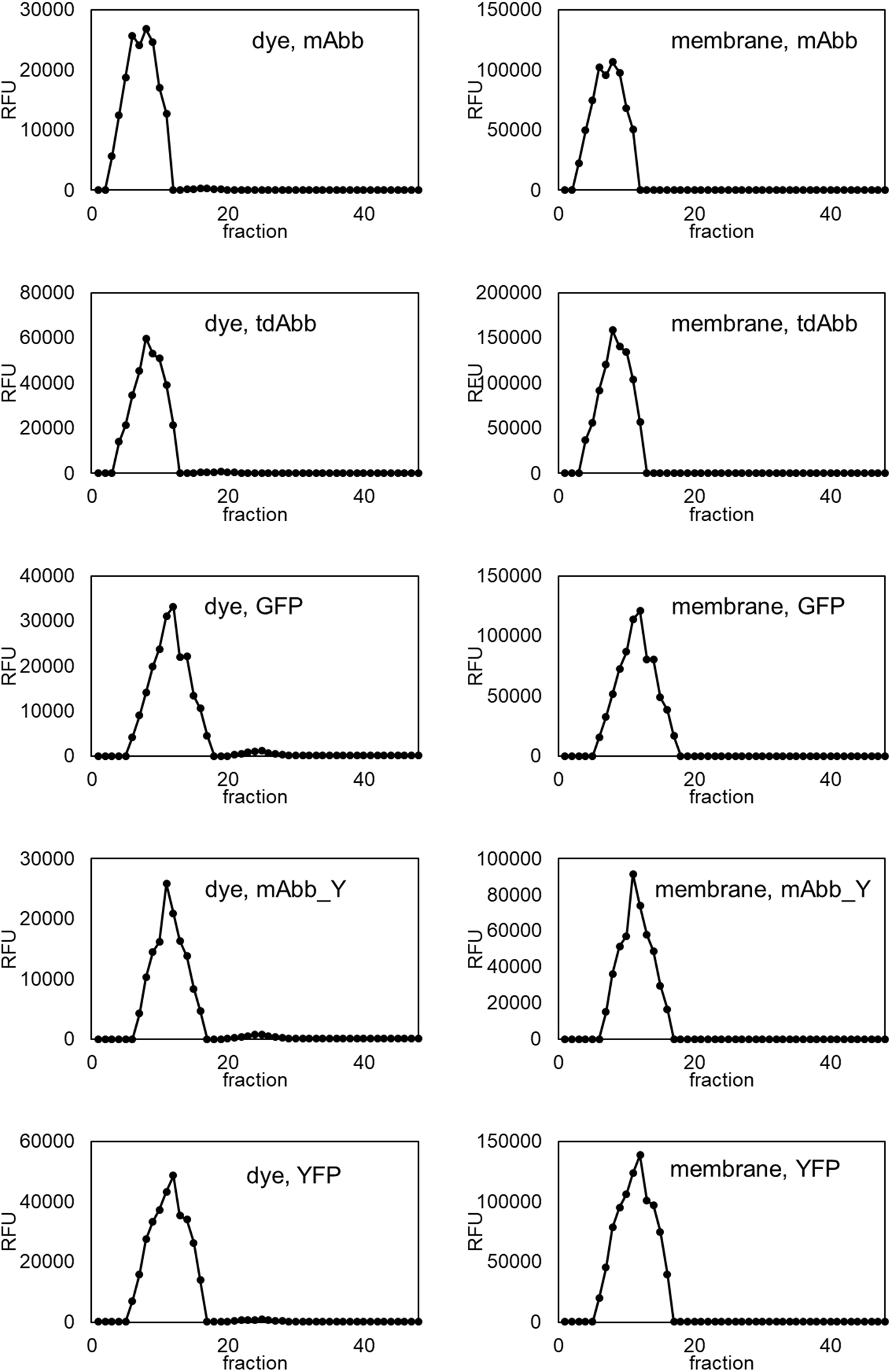
Representative liposome purification traces for calcein leakage from synthetic cells expressing one of the new fluorescent proteins or GFP or YFP controls. The dye trace shows calcein channel, the membrane trace shows Rhodamine membrane dye channel.

**Table S1.**
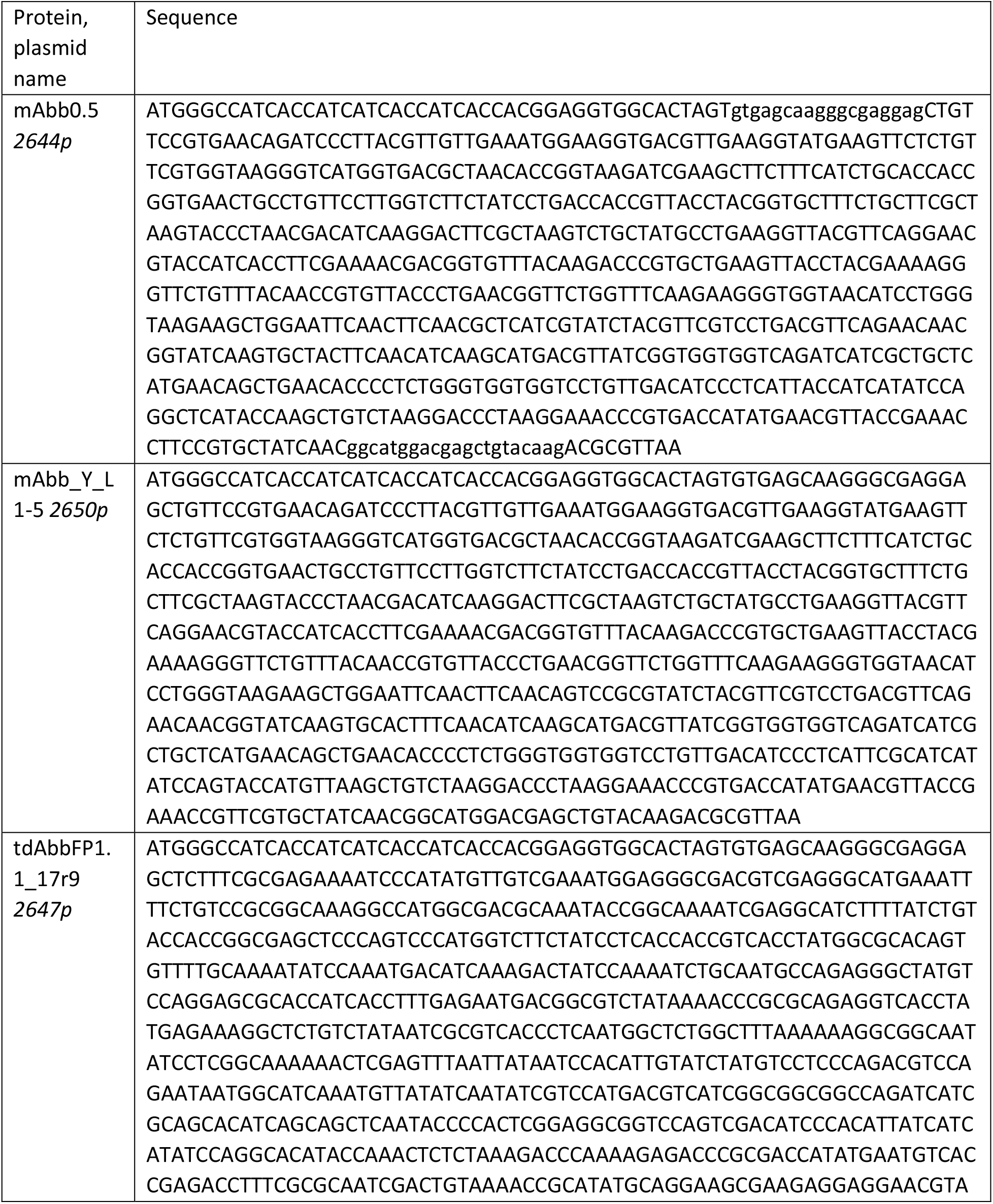
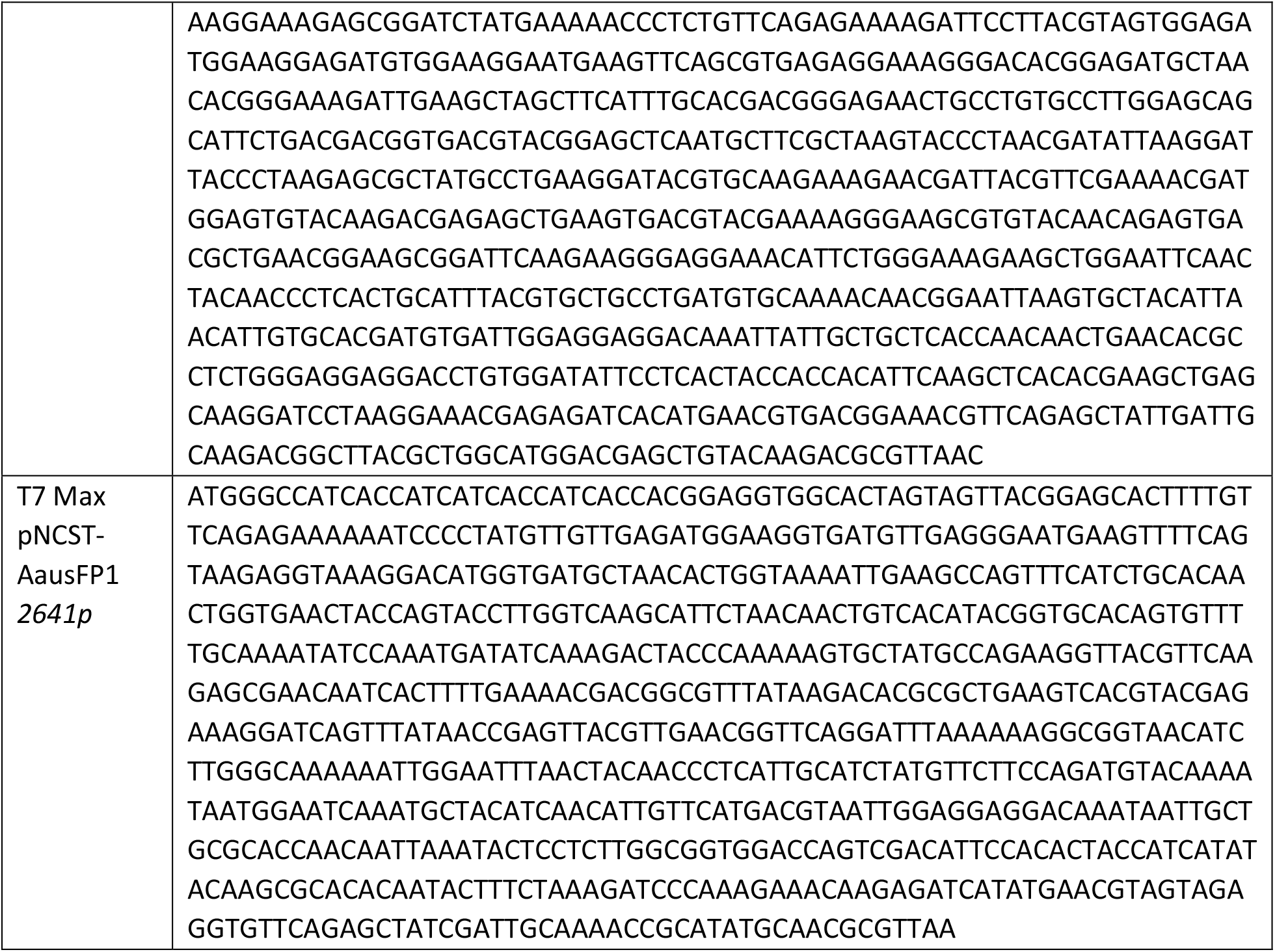
Proteins sequences

